# Behavioral anapyrexia as a response to virus infection in a poikilothermic vertebrate

**DOI:** 10.1101/2025.10.03.680298

**Authors:** Dávid Herczeg, Veronika Bókony, Gábor Herczeg, Dóra Holly, Andrea Kásler, János Ujszegi, Tibor Papp, Attila Hettyey

## Abstract

Behavioral plasticity may contribute to the ability of wild animals to survive disease outbreaks. In absence of endogen heat control, poikilothermic animals adjust their body temperature behaviorally. While many studies reported behavioral fever, its opposite, behavioral anapyrexia - when infected animals lower their body temperature by using cool microenvironments - remains poorly documented. Here, we report the first evidence of behavioral anapyrexia in a poikilothermic vertebrate as a response to pathogenic infection. We investigated thermoregulatory responses in tadpoles of the agile frog *Rana dalmatina*, a cool-adapted amphibian, following experimental infection with a warm-adapted ranavirus. Tadpoles were placed either in thermal gradients or homogeneously cool environments for five days post-exposure. In thermal gradients, all tadpoles reduced their preferred temperatures over time, but this decrease was steeper in infected tadpoles, and individuals with higher infection intensities preferred cooler temperatures. Infected tadpoles became increasingly precise in thermoregulation over time, while a similar trend was not detectable in non-infected tadpoles. Infection prevalence was similar between the two thermal environments, yet infection intensities were significantly higher in the thermal gradient. These results suggest fine-tuned thermoregulation by infected tadpoles to balance out the benefits of behavioral anapyrexia for fighting a warm-adapted pathogen versus the immune-suppressive and developmental costs of low temperatures.

## Introduction

Behavioral thermoregulation is the use of specific behaviors by a poikilothermic organism to regulate its body temperature, allowing it to maintain thermal balance within a range that supports optimal physiological performance ^1,2^. Accordingly, behavioral thermoregulation can also shape the outcome of host-pathogen interactions ^3^. Poikilothermic hosts can respond to pathogen presence by displaying behavioral fever ^4^, i.e., attaining higher body temperatures by using warmer microhabitats after pathogen exposure ^5,6^. This way animals can create conditions more favorable for themselves than for the pathogen thanks to an enhanced immune function or by taking advantage of a thermal optimum mismatch between the pathogen and its host ^5,7–11^. Infections can induce behavioral fever in poikilotherms ^5,6^. Diverse range of pathogens, including viruses, have been shown to trigger behavioral fever in amphibians ^12–15^, a taxon acutely threatened by disease pandemics and extinction risk worldwide ^16^.

Ranaviruses (hereafter *Rv*), belonging to the *Iridoviridae* family, evolved to infect a wide range of poikilothermic vertebrates, causing ranavirosis ^17^. Although ranavirosis alone cannot be held responsible for amphibian declines on a global scale, local epidemics have led to mass mortality events and, in some cases, resulted in the local extirpation of populations ^18–20^. Furthermore, the widespread distribution ^21^ and broad host range ^21,22^ of *Rv*s, coupled with their capability of interclass transmission between poikilothermic vertebrates ^23,24^, as well as the devastating outcomes of co-infections involving several *Rv*s or an *Rv* along with pathogenic bacteria or fungi ^25^ altogether confer *Rv*s dangerous potential for inducing massive collapses of amphibian populations on a global scale.

Temperature is the abiotic factor with perhaps the largest influence on the disease dynamics of ranavirosis in amphibians ^26^. Attempts to determine the thermal optimum range of *Rv* delivered contradictory results. Under *in vitro* conditions, the *Rv* strain Frog Virus 3 (FV3) replicates successfully between 8 and 30 °C, with a lower replication rate below 15 °C and the highest at 30 °C ^27,28^. This thermal dependency of *Rv* propagation is supported by the observation that *Rv*-related mortality is most frequent during the warm summer months in wild amphibian populations ^17^. Additionally, increasing the air temperature from 20 to 27 °C was shown to enhance *Rv* propagation, disease incidence, and mortality rates in the common frog *Rana temporaria* ^26^. Also, *Rv*-infected common frog tadpoles suffered higher mortality at 20 °C compared to 15 °C ^29^. Similarly, tadpoles of four amphibian species exposed to FV3 at 25 °C were consistently more likely to become infected and die than conspecifics exposed at 10 °C ^30^. In contrast, *Rv* infection probability and mortality were lower at 22 compared to 14°C in two *Lithobates* frog species infected with three different FV3 strains ^31^. Moreover, in larvae of the agile frog *R. dalmatina*, exposure to 30 °C for six days resulted in reduced prevalence and infection intensity of FV3, accompanied by enhanced host survival, as compared to conspecifics exposed to either 28 or 22 °C ^32^. Finally, worth to note that even slight temperature changes can strongly influence the outcome of FV3 infections: a 2 °C increase (from 10 to 12 °C) in wood frog *Lithobates sylvaticus* larvae turned apparently persistent infections into lethal ^30^. While these studies together highlight the effect of temperature on the progression of ranavirosis, inconsistencies in the observed patterns may stem from interspecific differences in the temperature-dependence of amphibian immune functions, in the temperature optima of *Rv* strains, or from varied outcomes of interactions between pathogen genotype, host genotype, and temperature ^31^. It is important to point out that in all aforementioned studies, environmental temperatures were set by the researchers, so that the tested amphibians did not have the opportunity to exert their thermal preferences and adjust their body temperatures *via* behavioral thermoregulation. The only study we know of that provided *Rv*-infected amphibians with the opportunity to thermoregulate reported that metamorphic and adult *Anaxyrus terrestris* toads displayed behavioral fever. More specifically, metamorphic individuals exposed to *Rv* increased their preferred temperature by 3.52 °C one day post-exposure, while adults exhibited a 1.43 °C increase in preferred temperature two days post-exposure as compared to non-infected controls ^12^.

Here, we investigate whether larval agile frogs adjust their thermal preferences following exposure to *Rv*, and whether the opportunity for thermoregulation can reduce infection prevalence and intensity. To achieve these objectives, we placed *Rv*-exposed and *Rv*-unexposed tadpoles into homogeneously cool environment or into thermal gradients for five days, recorded their thermoregulatory behavior, and measured infection intensities at termination. This setup allowed us to (i) study the thermoregulatory behavior and temperature optimum of *Rv*-unexposed tadpoles, (ii) detect potential infection-induced changes in thermoregulatory behavior, (iii) relate infection intensities to the magnitude of adjustments in thermoregulatory behavior, (iv) observe potential changes in general movement activity in parallel to disease progression, and (v) assess the effects of a homogeneously cool *vs*. variable thermal environment on virus replication. Based on previous findings of lower *Rv* infection intensities coupled with higher survival at high temperatures in agile frog tadpoles ^32^, we expected behavioral fever in response to *Rv* infection. We predicted that infected tadpoles would prefer higher body temperatures than their healthy conspecifics in the thermal gradient, and that infected tadpoles with higher infection intensities will prefer higher temperatures. Additionally, we predicted activity of infected animals to decrease over time, with lethargy emerging as the disease progressed ^33^, and to observe a steeper decline in the homogeneously cool environment (where high activity does not deliver obvious benefits) than in the thermal gradient (where behavioral thermoregulation may require moving around). We also expected infection intensities to be higher in the homogeneously cool environment as compared to the thermal gradient that had higher mean temperature and provided opportunity for thermoregulation (i.e., to express behavioral fever).

## Results

Across the five days, tadpoles in all treatment groups combined selected relatively high temperatures in the thermal gradient (22.4 °C ± 0.12 SE), both compared to the homogeneously cool environment (17.5 °C ± 0.12 SE; t_137_ = 33.66, *P* < 0.001; Supplementary Fig. S1A) and also compared to the average of the temperatures available in the thermal gradients (see non-overlapping CIs in Fig. 1). The preferred body temperatures (T_pref_) of tadpoles in the thermal gradient was significantly affected by the two-way interaction of *Rv*-status and the number of days since exposure (Table 1, Fig. 1). The T_pref_ significantly declined in all treatment groups over time during the experiment (Supplementary Table S1), where the decrease was significantly steeper in the *Rv*^+^ group compared to the *Rv*^-^ group and the *Rv*^0^ group, while there was no difference between the latter two (Supplementary Table S2; Fig. 1). As a result, by the 5^th^ day, T_pref_ was significantly lower in the in *Rv*^+^ group compared to the other two groups, and already from the 3^rd^ day compared to the *Rv*^0^ group (Fig. 1). The T_pref_ of *Rv*^0^ and *Rv*^-^ groups were not significantly different on either day (all *P* > 0.367). Furthermore, we found a significant negative effect of *Rv* infection intensity on the T_pref_ of *Rv*^+^ tadpoles (Table 1) i.e., tadpoles with higher *Rv* infection intensity had lower T_pref_.

**Fig. 1.**
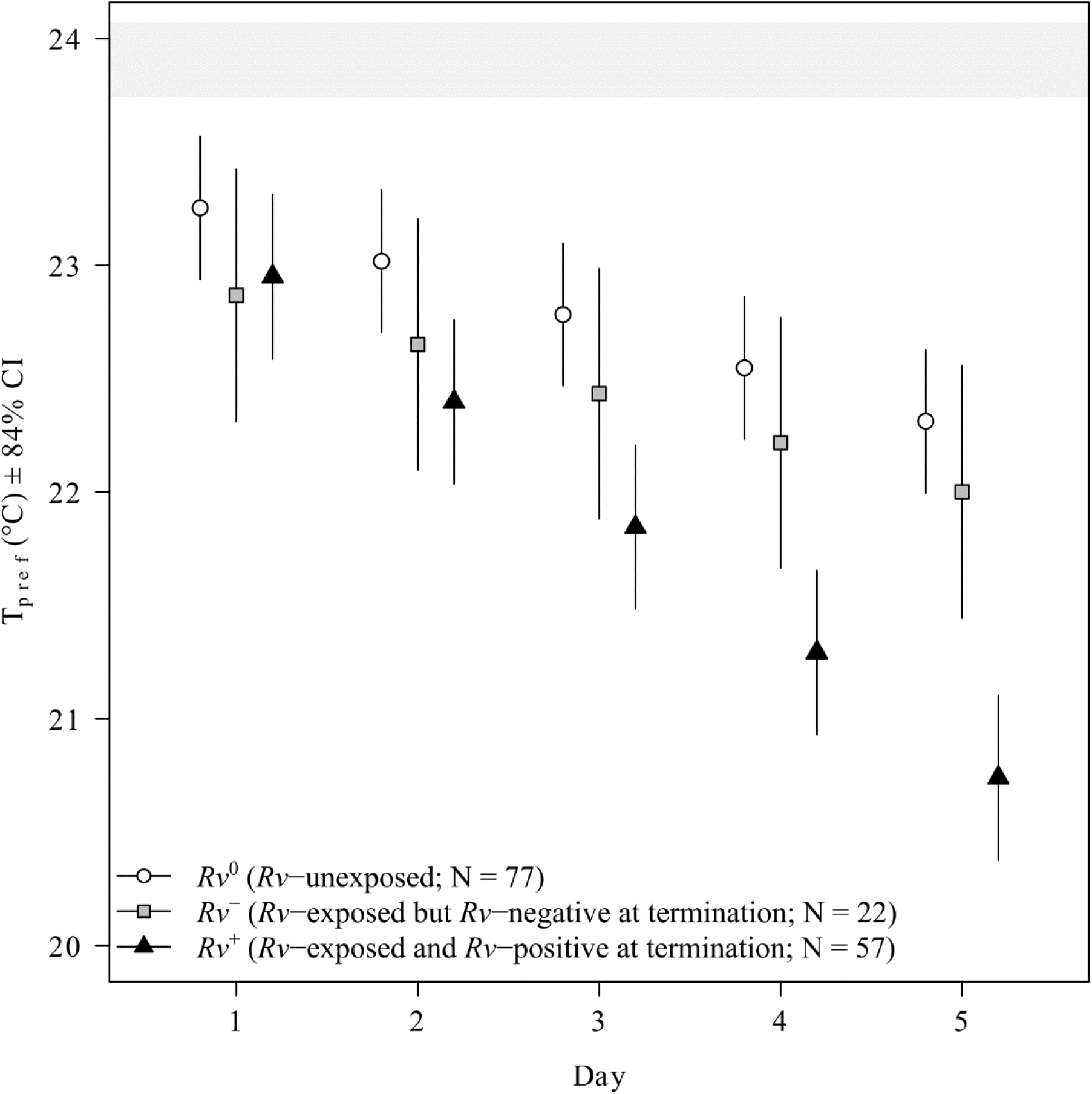
Preferred body temperatures (T_pref_) of agile frog tadpoles during the 5 days of the experiment. Means ± 84% Confidence Intervals (CIs) are shown. The horizontal grey area shows the 84% CI of the temperatures available in the experimental arenas, representing the body temperatures of a hypothetical thermoconformer tadpole that moves randomly in the thermal gradient. T_pref_ CIs that do not overlap with the grey area indicate active behavioral thermoregulation, i.e., non-random choice of thermal environment.

**Table 1.** Type-2 analysis-of-deviance tables of the statistical models. Significant effects (*P* < 0.05) are highlighted in bold.

| Dependent variable | Predictors | $\chi^2$ | df | $P$ |
| --- | --- | --- | --- | --- |
| Prevalence | Thermal environment | 0.006 | 1 | 0.935 |
|  | Round of experiment | 4.599 | 1 | **0.031** |
| Infection intensity | Thermal environment | 7.870 | 1 | **0.005** |
|  | Round of experiment | 0.028 | 1 | 0.865 |
| $T_{\text{pref}}$ | $Rv$-status | 9.465 | 2 | **0.008** |
|  | Day | 1092.26 | 1 | < **0.001** |
|  | Round of experiment | 4.315 | 1 | **0.037** |
| | $Rv$-status×Day | 216.452 | 2 | < **0.001** |
| $T_{\text{IQR}}$ | $Rv$-status | 2.271 | 2 | 0.321 |
|  | Day | 0.407 | 1 | 0.523 |
|  | Round of experiment | 4.612 | 1 | **0.031** |
| | $Rv$-status×Day | 6.181 | 2 | **0.045** |
| Activity | $Rv$-status | 1.850 | 2 | 0.396 |
|  | Day | 1.519 | 1 | 0.217 |
|  | Thermal environment | 1.030 | 1 | 0.310 |
| | $Rv$-status×Day | 229.938 | 2 | < **0.001** |
| | $Rv$-status×Thermal environment | 0.195 | 2 | 0.906 |
| | Day×Thermal environment | 9.756 | 1 | **0.002** |
| | $Rv$-status×Day×Thermal environment | 22.756 | 2 | < **0.001** |
| $T_{\text{pref}}$ on Day 5 in $Rv^+$ individuals | $\log(Rv)$ | 8.448 | 1 | **0.003** |
|  | Round of experiment | 3.152 | 1 | 0.075 |
| Activity on Day 5 in $Rv^+$ individuals | $\log(Rv)$ | 1.712 | 1 | 0.190 |
|  | Thermal environment | 1.167 | 1 | 0.682 |
|  | Round of experiment | 0.669 | 1 | 0.413 |
| | $\log(Rv)\times$Thermal environment | 2.869 | 1 | 0.090 |

$T_{\text{pref}}$ = Preferred body temperature; $T_{\text{IQR}}$ = Interquartile range of temperatures selected over a day; $Rv$ -status = Infection status on day 5 ( $Rv^0$ : $Rv$ -unexposed; $Rv^-$ : $Rv$ -exposed but $Rv$ -negative at termination; $Rv^+$ : $Rv$ -exposed and $Rv$ -positive at termination); Day = The number of days since exposure; $\log(Rv)$ = natural log-transformed ranavirus infection intensity

The interquartile range of temperatures selected over a day (T_IQR_) of tadpoles in the thermal gradient was significantly affected by the two-way interaction of *Rv*-status and the number of days since exposure (Table 1; Fig. 2). Specifically, T_IQR_ decreased over time in the *Rv*^+^ group, but not in the other two groups (Supplementary Table S1). However, there was no overall difference in the extent of T_IQR_ according to *Rv*-status (Supplementary Table S2), nor was there a significant difference on any one day (all *P* > 0.356).

**Fig. 2.**
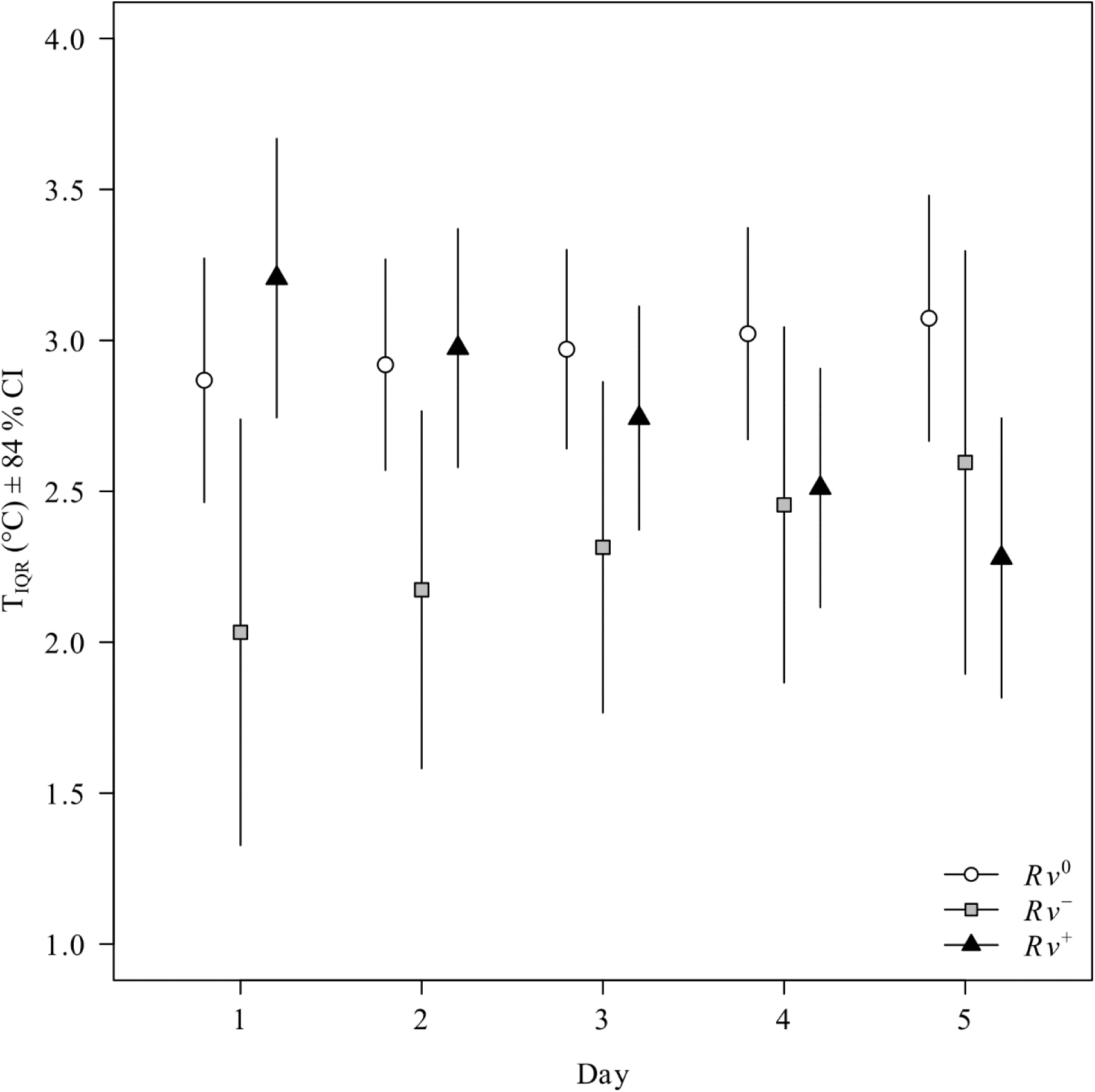
The interquartile range of temperatures selected over a day (T_IQR_) by agile frog tadpoles in thermal gradients.

Five days post-exposure, 72.2 % of *Rv*-exposed individuals had detectable *Rv* loads (*Rv*^+^) whereas 27.8 % were *Rv*-negative (*Rv*^−^). Amongst the *Rv-*exposed individuals, infection prevalence did not differ between the thermal gradient (75.6 %) and the homogeneously cool environment (76.5 %, Table 1). However, infection intensity in *Rv*^+^ individuals was significantly higher in the thermal gradient (9.42 × 10^7^ pfu ml^−1^ ± 3.96 × 10^7^ SE) than in the homogeneously cool environment (2.79 × 10^6^ pfu ml^−1^ ± 1.12 × 10^6^ SE; incidence rate ratio: 33.7 ± 19.5 SE, z = 6.08, *P* < 0.001; Supplementary Fig. S1B).

The movement activity of tadpoles was significantly influenced by the three-way interaction of the type of thermal environment, *Rv*-status and the number of days since exposure (Table 1; Supplementary Table S2; Fig. 3). The temporal change of activity varied across all treatment combinations except for *Rv*^0^ individuals between the two thermal environments (Supplementary Table S2, Fig. 3). Specifically, *Rv*^0^ tadpoles showed a slight increase in activity over time, but this was significant only in the thermal gradient; *Rv*^+^ tadpoles decreased activity with time in both environments, with a steeper decrease in the homogeneously cool environment; and *Rv*^−^ tadpoles increased activity with time in both environments, with a steeper increase in the homogeneously cool environment (Supplementary Table S2, Fig. 3). We found no effect of infection intensity on activity patterns on day 5 (Table 1).

**Fig. 3.**
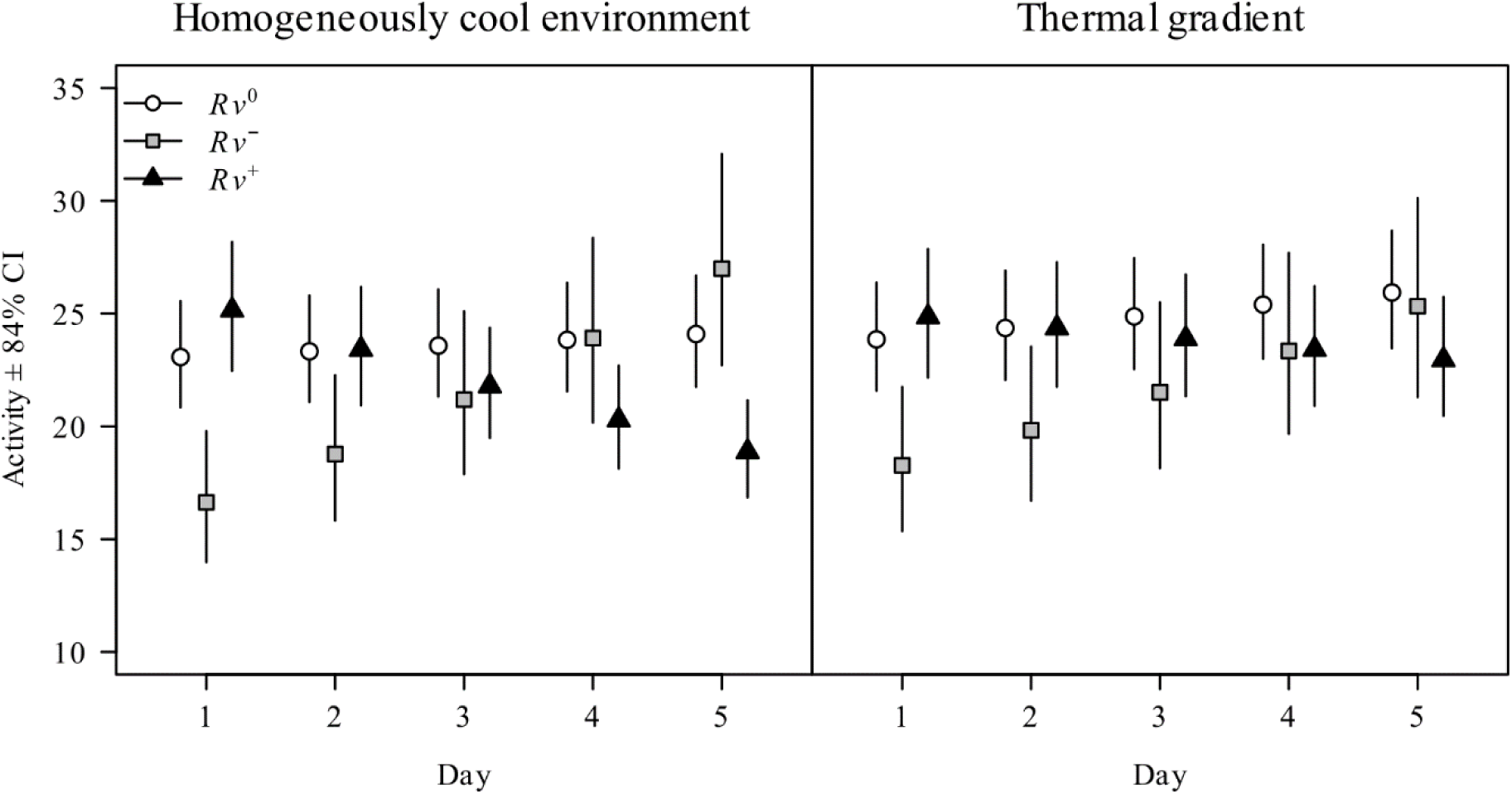
Activity of agile frog tadpoles during the experiment. Activity was estimated by measuring the distances between consecutive positions of tadpoles determined from recordings taken 2 min apart.

## Discussion

The key finding of this study was that in the thermal gradient, infected tadpoles showed lower preferred body temperatures than non-infected conspecifics already a few days after infection. Simultaneously, infected tadpoles thermoregulated with increasing accuracy over time, while this change was not observable in healthy tadpoles. After 5 days of disease progression, infected tadpoles with higher infection intensity had lower preferred body temperatures. Infection intensity was close to two magnitudes higher in thermal gradients than in homogeneously cool environments, as tadpoles in the latter were kept at lower temperatures compared to the temperatures selected by tadpoles in thermal gradients. Finally, we detected decreasing activity in *Rv*-positive individuals in both environments, but the pattern was stronger in the homogeneously cool environment.

Adjustments in thermoregulatory behavior in response to pathogens include behavioral fever, which has been documented in various invertebrates and poikilothermic vertebrates, including amphibians ^7,8,13–15,34–37^, even if its importance is debated, and the frequency of its occurrence across taxa is unknown ^5,38^. At the same time, an *in vivo* experiment suggested that exposure to high temperatures of around 30 °C can benefit agile frog tadpoles by reducing the severity of *Rv* infection ^32^. Consequently, we expected agile frog tadpoles to show behavioral fever. Contrary to our predictions and previous results, we found that *Rv*-infected agile frog tadpoles displayed behavioral anapyrexia from the third day after exposure to *Rv*. It is important to note that in the thermal gradient all animals displayed active thermoregulatory behavior, as both unexposed and infected tadpoles spent most time at temperatures that were on average lower than the mean temperature of their thermal environment. Also, individuals with higher infection intensities displayed stronger behavioral anapyrexia. At the same time, tadpoles in the homogeneously cool environment exhibited lower infection intensities than those in thermal gradients, although temperatures provided by the former could also be reached in the latter. Consequently, infected tadpoles displayed behavioral anapyrexia by lowering their thermal preference compared to non-infected tadpoles, but not to the extent to maximize the anti-pathogen effect. This discrepancy may be explained by the contrasting needs and constraints for optimizing body temperature under *Rv* infection. While the immune system of the poikilothermic hosts may work more efficiently at higher temperatures ^39,40^, this beneficial effect may be offset by higher virus replication rates ^26,30^ or by other costs associated with higher temperatures. Specifically, agile frog larvae experiencing high temperatures (28-30 °C) during larval development can suffer from increased mortality, delayed metamorphosis (without resulting in mass gain), and sex reversal ^41^. Consequently, behavioral fever expressed upon infection with *Rv* may not pay off for this cool-adapted host species. However, this does not explain the surfacing of behavioral anapyrexia.

Behavioral anapyrexia ^42,43^, previously referred to as ‘behavioral chill’ or ‘cold-seeking behavior’ was first documented in arthropods and was suggested to be a form of anti-pathogen behavioral defense ^44–47^. Similarly, the term ‘behavioral hypothermia’ has been reported in mice during bacterial infection ^48^ and in amphibians as a beneficial response to hypoxia, rather than as an anti-pathogen behavioral defense ^49^. The evolution of behavioral anapyrexia in agile frogs in response to FV3 may be explained by the warm-adapted nature of the virus, as *in vitro* studies have shown that FV3 reaches its highest titers at 30 °C ^27,28^. Consequently, low body temperature is likely to provide benefits to amphibian hosts because it presents unfavorable conditions for pathogen replication, while for anapyrexia to be advantageous to hosts also requires that lower temperatures cause a slighter decrease in their immune function than in *Rv* replication. The benefit of low temperatures is supported by some experimental studies performed *in vivo* on other host species ^26,29^. Considering that the agile frog is a cool-adapted species, this aligns well with the notion that the relationship between temperature and the effectiveness of the immune response may differ among warm- *vs.* cool-adapted amphibians: The former may benefit from higher temperatures, while the latter may do best under cooler conditions ^38,50,51^, even if this relationship can be further complicated by additional environmental factors ^11^. Whatever evolutionary force has led to its appearance; our study is the first to experimentally demonstrate pathogen-induced behavioral anapyrexia in a poikilothermic vertebrate.

Interestingly, infected tadpoles showing behavioral anapyrexia in the thermal gradients did not target as low body temperatures as what their conspecifics experienced in the homogeneously cool environments and what resulted in lower infection intensity, even though such temperatures (and even lower) were available in the gradient. One explanation may be that their target of thermoregulation shifted due to a constant reassessment of the trade-off between the anti-pathogen benefits and the physiological costs of maintaining body temperatures below the optimum. Such suboptimal temperatures can reduce immune function ^51,52^, metabolism, and locomotion^53,54^, and limit behaviors such as feeding, reproduction, and predator avoidance ^54,55^. The temporal dynamics of *Rv* infection is heavily influenced by the host’s immune system, which is largely shaped by ambient temperature during pathogen exposure and the infection dose ^30,32,33,56,57^. For example, American bullfrog *Lithobates catesbeianus* tadpoles experienced exponential growth of *Rv* titres within 2 days post-exposure, with replication rates rapidly increasing by 1-week post-exposure, a point at which tadpole mortality began to occur ^57^. It is also possible that if our experiment had lasted longer, infection intensities would have increased further, forcing infected tadpoles to choose even lower temperatures, further closing the gap between the temperature experienced in the homogeneously cool environments and T_pref_ in the thermal gradients. This is supported by the observation that T_pref_ decreased constantly from day to day and this curve has not yet levelled out by the 5^th^ day.

The determination of preferred body temperature, i.e., the body temperature voluntarily chosen in an ecological cost-free thermal gradient, is of central importance for understanding behavioral thermoregulation ^58^. Studies exploring changes in behavioral thermoregulation as anti-pathogen behaviors focused so far on mean temperatures ^12–15^, even though thermal biology considers the ranges of selected body temperatures to be of the essence ^58^. This is because thermoregulating animals typically do not aim to maintain one exact temperature value, but to keep their body temperature within a range. Therefore, approaching induced shifts in the target *via* estimating solely the mean, without considering the width of the range, is of limited use. Here, by showing that infected tadpoles did not only decrease their target body temperature but also increased their thermoregulatory precision, we demonstrate another layer of active behavioral anapyrexia, suggesting that the importance of accurate thermoregulation increased following infection. Thus, we encourage future studies on thermoregulatory responses to pathogens to incorporate both T_pref_ and T_IQR_ in their methodological framework.

According to our predictions, activity decreased over time in *Rv*^+^ tadpoles in both the homogeneously cool environment and the thermal gradient, with a steeper decrease observed in the former. This decrease in activity might be explained by an automatic consequence of lower preferred temperatures, as activity also declined in the homogenously cool environment where the temperature remained stable. However, *Rv*-induced activity decrease was not detected within a comparable timeframe to ours in post-metamorphic individuals of either wood frogs *Lithobates sylvaticus* ^59^ or agile frogs ^60^, and has so far only been detected in the terminal stage of disease progression in tiger salamanders *Ambystoma tigrinum* ^61^. In our case, the observed activity decreases in *Rv*^+^ tadpoles may have resulted from accumulating pathological damage or the energetic costs associated with an intensifying immune response during disease progression - most likely a combination of both. At constant lower temperatures applied in our experiment, immune responses may be less effective in amphibians ^40^, requiring more sustained metabolic effort to fight infections, thereby leaving less energy available for locomotion ^62^. In contrast, within the thermal gradient, infected tadpoles may have used thermoregulation to optimize energy allocation more efficiently, allowing them to maintain higher levels of activity.

Curiously, we found increasing activity with time in the *Rv^−^*tadpoles, both in the homogeneously cool environment and the thermal gradient, with a steeper increase observed in the former. This increase started from a low level of activity which gradually reached the level observed in the *Rv*^0^ group. It is unclear if in *Rv^−^* tadpoles the experimental infection was successful but these individuals cleared *Rv* in five days or if experimental exposure to *Rv* was unsuccessful in this group of animals. The observed low activity immediately following exposure to *Rv* supports the former scenario. We speculate that these individuals exhibited a stronger/faster than average immune response to the pathogen, which in turn decreased their movement activity in the beginning. Ascertaining this hypothesis would, however, require targeted investigation.

In conclusion, we detected behavioral anapyrexia in an amphibian - virus host-pathogen system, where infected animals lowered their preferred temperatures by around 1.6°C by the 5^th^ day post-infection as compared to non-infected conspecifics. The strength of the behavioral response of agile frogs depended on infection intensity. Moreover, individuals maintained in a thermal gradient had roughly twice as high infection intensity than conspecifics maintained at the homogenously cool environment. This suggests that, at least within the 21–23 °C range, maintaining a lower body temperature in the presence of *Rv* may be advantageous to combat the infection. These two results together suggest that behavioral anapyrexia might be an adaptive response to infection with *Rv*, but further experimental investigations will be necessary to prove this. Our observation that infected tadpoles decreased their target body temperature and at the same time increased their thermoregulatory precision draws attention to the fact that studies concerned with disease-induced changes in thermoregulatory behavior should not only focus on the widely used T_pref_ but should also assess T_IQR_ to achieve a deeper understanding. In addition, we detected decreasing activity in *Rv*-infected tadpoles, which we interpret as a sign of progressing disease. We note that a longer observation period could have provided further information on how far behavioral anapyrexia, infection intensity and decreasing activity would have progressed, However, with longer observation periods, it becomes difficult to disentangle whether behavior shapes infection status or, conversely, infection shapes behavior. We speculate that both mechanisms act reciprocally. Under natural circumstances, behavioral anapyrexia may manifest in seeking shaded areas and deeper water, or daily activity shifted towards cooler parts of the day. Such behavioral changes may, however, pose fitness costs, which may be exacerbated by intensifying climate change. Our results suggest that infected tadpoles engage in fine-tuned thermoregulation to balance the benefits of behavioral anapyrexia against a warm-adapted pathogen with the immune-suppressive and developmental costs of lower body temperatures. Understanding how temperature simultaneously affects pathogens and host physiology is key to clarifying host–pathogen dynamics and guiding amphibian conservation.

## Materials and methods

All experimental procedures were approved by the Ethical Commission of the Plant Protection Institute, and permissions were issued by the Government Agency of Pest County (PE-06/KTF/8060-1/2018, PE-06/KTF/8060-2/2018, PE-06/KTF/8060-3/2018, PE-06/KTF/8060-5/2018, PE/EA/295-7/2018 and PE/EA/58-4/2019). The experiments were carried out according to recommendations of the EC Directive 86/609/EEC for animal experiments (http://europa.eu.int/scadplus/leg/en/s23000.htm).

### The study species

The agile frog (*Rana dalmatina* Fitzinger, 1838) is endemic to large parts of continental Europe and southern Scandinavia. It occupies a wide variety of aquatic and terrestrial habitats at altitudes ranging between zero to 1700 m a.s.l. ^63^. Over much of its distribution range it is the first anuran amphibian to start yearly breeding activities, spawning already in late winter or early spring ^64,65^. The critical thermal maximum of agile frog tadpoles raised in the wild is 36.8 °C ^66^, which is relatively low compared to that of other temperate amphibian larvae ^67^. This indicates that the agile frog is a rather cool-adapted species.

### Animal collection and husbandry

In late March and early April 2021, we collected agile frog eggs on two occasions separated by one week. On both occasions we collected ca. 100 eggs from each of five freshly laid clutches (hereafter sib-groups) from two natural populations (Nagykovácsi Békás-tó 47°34’34“N, 18°52’08“E and Lendület-tó 47°33’04“N, 18°55’36“E) located in the vicinity of Budapest, Hungary. We transported eggs to the nearby Júliannamajor Experimental Station of the Plant Protection Institute. Until hatching, we reared embryos separated by sib-groups in plastic dishpans (24 × 16 × 13 cm) holding 1 liter of reconstituted soft water (RSW) ^68^. We maintained a temperature of 15.49 ± 0.92 °C (mean ± SD) and a 12:12 h light:dark cycle.

After hatching, each sib-group was haphazardly divided into three groups of 16 healthy larvae and placed into 15-L opaque, plastic rearing containers (37 × 27 × 16.5 cm), each holding 10 liters of RSW. We changed the RSW twice a week and fed tadpoles with chopped and slightly boiled spinach *ad libitum*. When individuals reached developmental stage 25 (Day 0) according to Gosner ^69^, we haphazardly selected 12 healthy tadpoles (i.e., we removed surplus individuals) from each rearing container and randomly assigned them to treatment groups (*Rv*-exposed *vs*. *Rv*-unexposed). Surplus individuals were released at their place of origin. On Day 10 we exposed half of the tadpoles individually to *Rv* (Frog Virus 3; ATCC No. VR-567) by placing them into plastic cups holding 500 ml RSW and *Rv* at a concentration of 3 × 10^4^ plaque forming units (pfu) × ml^−1^. Experimental exposure lasted for 24 hours. Simultaneously, we exposed the other half of tadpoles to a sham extract (i.e., *Rv*-free medium containing 2 % fetal bovine serum) following the same procedure. Maintenance of *Rv* cultures followed earlier protocol ^32^.

### Experimental trials

During the experiment, laboratory air temperature was 18.69 ± 1.67 °C (mean ± SD), and we maintained a 13:11 h light:dark cycle. Due to spatial limitations in the laboratory and to increase our sample size, we performed two rounds of experimental trials consecutively, corresponding to the two cohorts of animals collected as eggs one week apart. We made our observation with 80 individuals in each round, using each individual once. We set up ten metal trays (206 × 72 × 13.5 cm, length × width × height, respectively) on two shelves using five columns of a laboratory shelf system, and we filled trays with tap water (Supplementary Fig. S2A). Along the longitudinal axis, each tray was separated into four equal compartments by dividers made of 5 cm thick XPS insulation board. On top of the dividers, we fixed eight half-pipes (200 × 8 × 8 cm, length × width × height, respectively) and filled them with 4.8 liters of RSW, resulting in a water depth of ca. 3 cm (Supplementary Fig. S2B). We kept water levels in half-pipes low to minimize thermal stratification and conductive water circulation. Half-pipes were partially immersed into the surrounding water to allow for heat exchange between the water in the trays and the water in the half-pipes. Half-pipes in five trays positioned on the lower level of the shelf system (approx. 40 cm above the ground) provided homogeneously cool environments for the tadpoles, where the water temperature followed the air temperature in the laboratory. The temperatures available to tadpoles along half-pipes in the homogenously cool environment varied slightly around 18 °C (17.92 ± 0.5 °C; mean ± SD) and in the thermal gradient between 19.15 ± 1.45 °C at the cool end and 30.57 ± 1.06 °C at the warm end (for details see Supplementary Fig. S3). Temperatures within this range are known to influence the progression of ranavirosis ^26,29,32^ and encompass the thermal optimum for agile frog tadpoles (22–24 °C; Hettyey et al., unpublished data). Moreover, such water temperatures are commonly encountered by tadpoles in their natural habitats ^66^. To create temperature gradients in the half-pipes, we heated the external water in one end compartment of trays to ca. 40 °C with a 400 W towel dryer heating element equipped with a thermostat. In the neighboring compartment we raised the water temperature to 30 °C using a submersible 300 W aquarium heater (Tetra HT 300). In the third compartment, we did not modulate water temperature in trays. In the fourth compartment we cooled the water in trays to 12 °C by circulating it through external water chillers (a Hailea HC-500A and a Hailea HC-300A connected in series). To mix the water within tray compartments, we equipped each compartment with a water pump (Tetra WP 300). To minimize the dissipation of heat through the thin walls of trays and to ensure consistent temperature conditions in half-pipes throughout the experiment, we insulated the outside (i.e., the bottom and all four sides) of the trays with XPS insulation. The trays providing homogeneously cool thermal environments for tadpoles were set up in the same way as those providing thermal gradients, except that the former lacked heating and cooling equipment. Between rounds, the half-pipes were removed, emptied, cleaned and disinfected with 70 % ethanol to avoid cross-contamination. Half-pipes were assigned randomly to *Rv* treatments.

On Day 11, we measured the water temperature along the half-pipes using digital thermometers (Greisinger GTH 175/PT) at mid-depth of the water column. We took measurements at 20-cm intervals in the case of thermal gradients and at 50-cm intervals in the case of homogeneously cool environments. We performed temperature recordings twice before the first round and twice before the second round. After temperature measurements, we started experimental trials by placing one tadpole into the center of each half-pipe. Over the following five days (Days 12-16), we recorded the thermoregulatory behavior of tadpoles by taking video recordings (30 fps) for 12 sec, each followed by a 2 min break, for three hours each day between 10 am and 1 pm, using car cameras (Concorde RoadCam HD 10) positioned above the half-pipes. This resulted in 78 video recordings per individual for each of the 5 days of observation. Following the last video recording each day, we fed each tadpole with 5 food pellets (Sera GranuGreen) placed at 20 cm intervals along the half-pipes to provide *ad libitum* food at any one feeding spot and thereby avoid forcing hungry tadpoles into areas they would otherwise not move to. Uneaten food pellets and excrements of tadpoles were removed every morning before the start of the first video recording to avoid confounding effects of foraging behavior on thermoregulatory behavior. To compensate for evaporation, after food removal we topped up water to the original level in half-pipes with reverse-osmosis filtered water of 18 °C or pre-heated to 25 °C (in the half-pipes providing a homogeneously cool environment or a thermal gradient, respectively). We finished this procedure before 9 am to allow at least one hour for the temperature to set before starting recordings. At the end of each round (1^st^ round: Day 16 and 2^nd^ round: Day 21), we measured the body mass of tadpoles to the nearest 0.01 g using a laboratory scale (Ohaus PA114), euthanized them ^70^ and conserved each in 3 ml of 96 % ethanol.

From the middle frame of the 12 sec recordings, we determined the spatial position of tadpoles. This was facilitated by a scale (ticks every 10 cm) drawn along the longitudinal axis of half-pipes on the bordering insulation. We estimated the temperature of the immediate thermal environment of the animals by projecting their spatial position (estimated to the nearest cm) on half-pipe specific maps of water temperatures. By averaging measurements across the two rounds, we generated half-pipe–specific water temperature maps, assuming quadratic plane-curve changes in temperature between adjacent points. In accordance with previous similar studies ^71^ we assumed that the body temperature of a given tadpole corresponded well with the temperature of its immediate thermal environment, justified by the small size of the tadpoles and the resulting negligible thermal inertia. We used these estimated temperature values as proxy for the preferred body temperatures (T_pref_) of tadpoles that were allowed to thermoregulate, ‘and we calculated the interquartile range of temperatures selected over a day (T_IQR_; equivalent to the width of the set-point range or T_set_ in the thermal ecology literature ^1,2^). Activity was estimated by measuring the distances between consecutive positions of tadpoles determined from recordings taken 2 min apart.

### Molecular analysis

To assess *Rv* infection intensity, we excised the livers and homogenized them with a disposable pellet mixer (VWR, catalogue no. 47747-370). We extracted DNA with Wizard Genomic DNA Purification Kits (Promega, Madison, Wisconsin, USA) according to the manufacturer’s protocol. We stored extracted DNA at -20 °C until further analysis. We estimated *Rv* infection intensities by a qPCR assay targeting the major capsid protein (MCP) gene of the virus following standard amplification methodologies ^72^. Dilutions of purified MCP amplicons were used for setting the standard curves and these had been correlated to infective virion titers in the same qPCR as detailed earlier ^32^. Reactions were run on a BioRad CFX96 Touch Real-Time PCR System.

### Statistical analysis

We created three categories of experimental animals according to their *Rv*-status, based on combinations of *Rv*-exposure history and infection status at the end of the experiment: (1) *Rv*-exposed and *Rv*-positive at termination (*Rv*^+^ henceforth), (2) *Rv*-exposed, but *Rv*-negative at termination (*Rv*^-^ henceforth), and (3) not exposed to *Rv* (*Rv*^0^ henceforth). We treated *Rv*^-^ individuals as a separate group because it remains unclear whether these *Rv*^-^ individuals were not successfully infected or if they cleared the infection during the experiment.

Mortality during the experiment was very low, with only a single *Rv*^0^ tadpole dying in the first round and a single *Rv*-exposed tadpole dying in the second round of trials. We found two individuals in the *Rv*^0^ group that were cross-contaminated with the virus, but infection intensities were very low in these cases (18 and 48 pfu ml^−1^, respectively). These individuals were excluded from the analyses. This resulted in the following sample sizes: 38 *Rv*^0^, 11 *Rv*^-^, and 29 *Rv*^+^ animals in the homogeneously cool environment and 39 *Rv*^0^, 11 *Rv*^-^, and 28 *Rv*^+^ animals in the thermal gradients.

First, we analyzed the relationship of T_pref_ and T_IQR_ with *Rv*-status focusing on tadpoles kept in the thermal gradients. To estimate T_pref_, we entered all temperature records of each individual as repeated measures in a linear mixed model (LMM). For estimating T_IQR_, we calculated the interquartile range of selected temperatures for each of the 5 days for each tadpole, and used these 5 values as repeated measures in an LMM. In both models, we included the round of the experiment (i.e., first *vs*. second round) as a fixed factor, while individual ID nested in sib-group and as well as half-pipe ID were entered as random factors. To allow for differences in the slope of temporal change of thermoregulatory behavior among the three treatment groups, in both models we included the interaction between *Rv*-status (fixed factor) and the number of days since exposure treated as a covariate.

Second, we analyzed the activity of all experimental animals by entering the distances between consecutive positions of each tadpole as repeated measures in a generalized linear mixed model (GLMM) with negative-binomial error. As fixed effects, the model included the type of thermal environment (i.e., homogeneously cool *vs*. gradient), *Rv*-status, the number of days since exposure, all their two-way interactions and their three-way interaction, and the round of the experiment. As random factors, we used the same random structure as for T_pref_ and T_IQR_.

Third, because *Rv* infection intensity was measured on the 5^th^ day of observation, we used the behavioral data of that day to analyze the effects of infection intensity on T_pref_ and activity within *Rv*^+^ individuals. These models had the same structure as above, but the fixed effects were infection intensity (ln-transformed for better model fit), round of the experiment, thermal environment and its interaction with infection intensity.

To compare the temperatures selected by tadpoles between thermal gradients and homogeneously cool environments, we used an LMM including the round of experiment as an additional fixed factor, and individual nested in sib-group as random factors. We allowed for different variance in the two types of thermal environment. Third, we tested the effects of the thermal environment on *Rv* prevalence, as well as on infection intensity among *Rv*+ using GLMMs, with binomial and negative-binomial error, respectively. The type of thermal environment and the round of the experiment were included as fixed factors, while half-pipe ID and sib-group were used as crossed random factors.

All analyses were run in R 4.3.2 ^73^, using the packages *’lme4*’ ^74^ (functions ‘*glmer*’ and ‘*lmer*’) for binomial and linear models, ‘*nlme*’ (‘*lme*’ and ‘*varIdent*’ functions) for the linear model with group-specific variances, ‘*glmmTMB*’ ^75^ for negative-binomial models, ‘car’ (function ‘*Anova*’) for analysis-of-deviance tables, ‘*emmeans*’ ^76^ for calculating marginal means and linear contrasts, and ‘*DHARMa*’ ^77^ for residual diagnostics. We used the false discovery rate (FDR) method to correct p-values for multiple comparisons across *Rv*-status groups and days ^78^. To illustrate significant differences in graphs, we present the model-derived mean estimates with 84% Confidence Intervals (CIs), because the lack of overlap between two such intervals is equivalent to a significant difference at 5% significance level ^79^.

## Data availability statement

Data and the annotated R script used for the statistical analysis is available from Figshare Digital Repository doi: 10.6084/m9.figshare.30273211. The DOI becomes active upon acceptance of the manuscript. For reviewers: please see the following private link to the data and code: https://figshare.com/s/93a2e2bb6e3e31bbc44c

## Acknowledgement

We thank Márk Szederkényi for field and technical assistance and for Zsófia Boros, Csenge Kalina, Réka Bertalan and Nikolett Ujhegyi for their help in conducting experiments. The authors are indebted to Krisztina Ursu (NÉBIH) and Márk Z. Németh (HUN-REN ATK NÖVI) for their technical assistance during laboratory work. We thank Andor Doszpoly (HUN-REN ÁTKI) for help with *Ranavirus* maintenance. The original Frog Virus 3 isolate was obtained from Rachel E. Marschang (University of Hohenheim, Stuttgart, currently at Laboklin GmbH). DHe and AH were supported by the János Bolyai Research Scholarship of the Hungarian Academy of Sciences. The study was performed with financial support from the National Research, Development and Innovation Office of Hungary (NKFIH; K-124375 and K-147500 to AH, PD-142654 to JU and STARTING-149841 to DHe) and the University Research Fellowship of the Ministry for Innovation and Technology (EKÖP-24-4 for JU). The authors were further supported by the New National Excellence Program of the Ministry for Culture and Innovation from the source of the National Research, Development and Innovation Fund (ÚNKP-20-3 and ÚNKP-21-3 to A.K.). This project has received funding from the HUN-REN Hungarian Research Network.

## Authors contributions

Dávid Herczeg and Attila Hettyey conceived the study and designed the methodology; Dávid Herczeg, Dóra Holly, Andrea Kásler, János Ujszegi, Tibor Papp and Attila Hettyey collected the data; Veronika Bókony analysed the data; Dávid Herczeg, Gábor Herczeg, Veronika Bókony and Attila Hettyey led the writing of the manuscript. All authors contributed critically to the drafts and gave final approval for publication.

## Competing interests

The authors declare no conflict of interest.

## Materials & Correspondence

Dávid Herczeg

## Notes

https://figshare.com/account/articles/30273211

### Competing Interest Statement

The authors have declared no competing interest.

