## Supplementary Material for "Behavioral anapyrexia as a response to virus infection in a poikilothermic vertebrate"

**Supplementary Table S1.** Behavioral changes with time (per day over 5 days) in each treatment group. Estimated marginal means of linear trends (slopes) from statistical models, associated standard errors (SE) and lower (LCL) and upper (UCL) 95% confidence limits, in each infection-status group are shown for temperature preference ($T_{pref}$) and interquartile range of selected temperatures over a day ($T_{IQR}$) in thermal gradients, and activity in both thermal environments. Slopes highlighted in bold differ from zero at a 5% significance level. For comparisons of slopes to each other, see Supplementary Table S2.

| **Dependent variable** | **Slope** | **SE** | **LCL** | **UCL** |
|---|---|---|---|---|
| **$T_{pref}$ in thermal gradient** | | | | |
| $Rv^0$ | **-0.235** | 0.0149 | -0.264 | -0.206 |
| $Rv^-$ | **-0.217** | 0.0278 | -0.271 | -0.162 |
| $Rv^+$ | **-0.553** | 0.0175 | -0.587 | -0.518 |
| **$T_{IQR}$ in thermal gradient** | | | | |
| $Rv^0$ | 0.051 | 0.085 | -0.116 | 0.219 |
| $Rv^-$ | 0.140 | 0.157 | -0.170 | 0.451 |
| $Rv^+$ | **-0.231** | 0.099 | -0.428 | -0.036 |
| **Activity in homogeneously cool environment** | | | | |
| $Rv^0$ | 0.010 | 0.006 | -0.001 | 0.022 |
| $Rv^-$ | 0.121 | 0.012 | 0.097 | 0.144 |
| $Rv^+$ | **-0.071** | 0.007 | -0.085 | -0.057 |
| **Activity in thermal gradient** | | | | |
| $Rv^0$ | 0.02 | 0.006 | 0.008 | 0.032 |
| $Rv^-$ | 0.081 | 0.012 | 0.057 | 0.106 |
| $Rv^+$ | **-0.019** | 0.007 | -0.033 | -0.005 |

$Rv^0$: *Rv*-unexposed; $Rv^-$: *Rv*-exposed but *Rv*-negative; $Rv^+$: *Rv*-exposed and *Rv*-positive

**Supplementary Table S2.** Pairwise comparisons of the slopes of behavioral changes over time. The infection-status groups are compared to each other in the thermal gradient for temperature preference ($T_{pref}$) and the interquartile range of selected temperatures over a day ($T_{IQR}$). Additionally, pairwise comparisons of infection-status groups for activity levels in both thermal environments are provided. Linear contrasts (*c*), associated standard errors (SE), and *P*-values adjusted using the FDR method are reported. Significant differences (P < 0.05) are in bold.

| Dependent variable | Pairwise comparison | *c* | SE | *P* |
|---|---|---|---|---|
| **$T_{pref}$ in thermal gradient** | | | | |
| | $Rv^0$ *vs*. $Rv^-$ | -0.018 | 0.031 | 0.559 |
| | $Rv^0$ vs. $Rv^+$ | 0.317 | 0.023 | < **0.001** |
| | $Rv^-$ vs. $Rv^+$ | 0.335 | 0.032 | < **0.001** |
| **$T_{IQR}$ in thermal gradient** | | | | |
| | $Rv^0$ *vs*. $Rv^-$ | -0.089 | 0.179 | 0.618 |
| | $Rv^0$ *vs*. $Rv^+$ | 0.283 | 0.131 | 0.069 |
| | $Rv^-$ *vs*. $Rv^+$ | 0.372 | 0.186 | 0.069 |
| **Activity** | $Rv^0$ hom. *vs*. $Rv^-$ hom. | -0.110 | 0.013 | < **0.001** |
| | $Rv^0$ hom. *vs*. $Rv^+$ hom. | 0.082 | 0.009 | < **0.001** |
| | $Rv^0$ hom. *vs*. *Rv*- gradient | -0.01 | 0.008 | 0.249 |
| | $Rv^0$ hom. *vs*. $Rv^-$ gradient | -0.07 | 0.013 | < **0.001** |
| | $Rv^0$ hom. *vs*. $Rv^+$ gradient | 0.03 | 0.009 | **0.001** |
| | $Rv^-$ hom. *vs*. $Rv^+$ hom. | 0.192 | 0.014 | < **0.001** |
| | $Rv^-$ hom. *vs*. $Rv^0$ gradient | 0.1 | 0.013 | < **0.001** |
| | $Rv^-$ hom. *vs*. $Rv^-$ gradient | 0.039 | 0.017 | **0.025** |
| | $Rv^-$ hom. *vs*. $Rv^+$ gradient | 0.14 | 0.014 | < **0.001** |
| | $Rv^+$ hom. *vs*. $Rv^0$ gradient | -0.092 | 0.009 | < **0.001** |

| | | | |
|---|---|---|---|
| $Rv^{+}$ hom. *vs*. $Rv^{-}$ gradient | -0.153 | 0.014 | **< 0.001** |
| $Rv^{+}$ hom. *vs*. $Rv^{+}$ gradient | -0.051 | 0.01 | **< 0.001** |
| $Rv^{0}$ gradient *vs*. $Rv^{-}$ gradient | -0.06 | 0.013 | **< 0.001** |
| $Rv^{0}$ gradient *vs*. $Rv^{+}$ gradient | 0.04 | 0.009 | **< 0.001** |
| $Rv^{-}$ gradient *vs*. $Rv^{+}$ gradient | 0.1 | 0.014 | **< 0.001** |

$Rv^{0}$: *Rv*-unexposed; $Rv^{-}$: *R*v-exposed but *Rv*-negative; $Rv^{+}$: *Rv*-exposed and *Rv*-positive; hom: Homogenously cool environment; gradient: thermal gradient

**Supplementary Figure S1.** Distribution of A) tadpole temperatures and B) *Rv* infection intensities in the two thermal environments. In each violin plot, the black dot and the white box represent the median and interquartile range, whiskers extend to 1.5 × interquartile range, and the grey polygon is a kernel density plot.

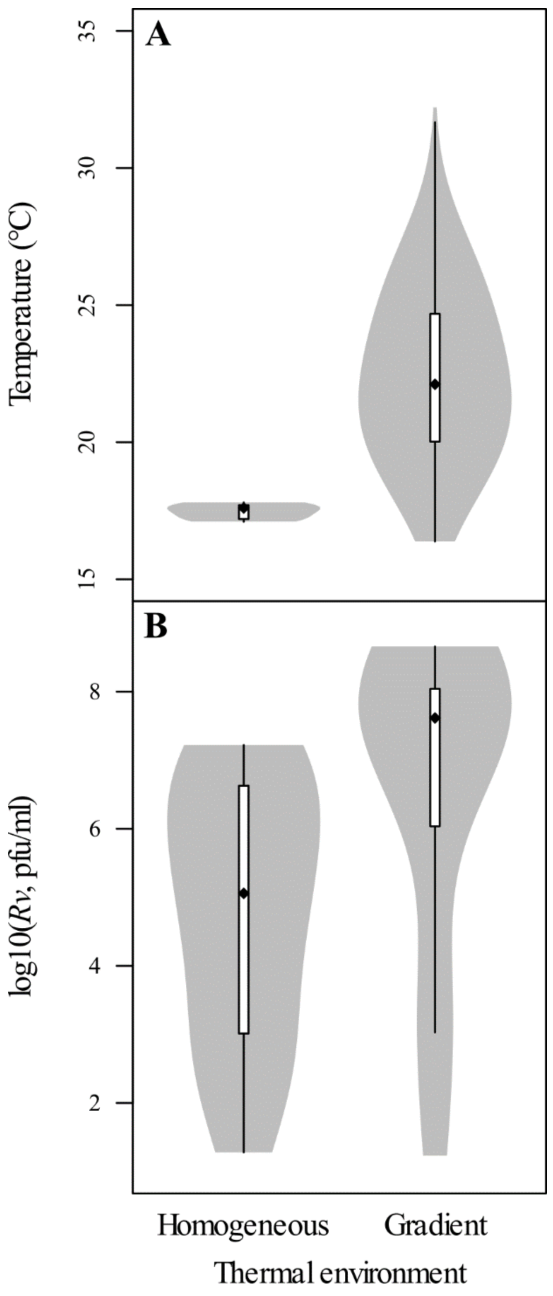

**Supplementary Figure S2**

The setups for providing different thermal environments in this study. (A) Half-pipes located on the upper level of the shelf system, creating thermal gradients and (B) half-pipes positioned on the lower level, providing homogeneously cool environments. Components: 1 – 400 W towel dryer heating element equipped with a thermostat; 2 – Warm end of insulated metal tray; 3 – Warm end of half-pipes; 4 – One of three dividers forming four compartments in the metal tray; 5 – Cool end of half-pipes; 6 – Tubes for cold water circulation (supplied by external water chillers) at the cool end of the tray; 7 – Car cameras used for behavioral recording; 8 – Agile frog tadpole in the homogeneously cool environment.

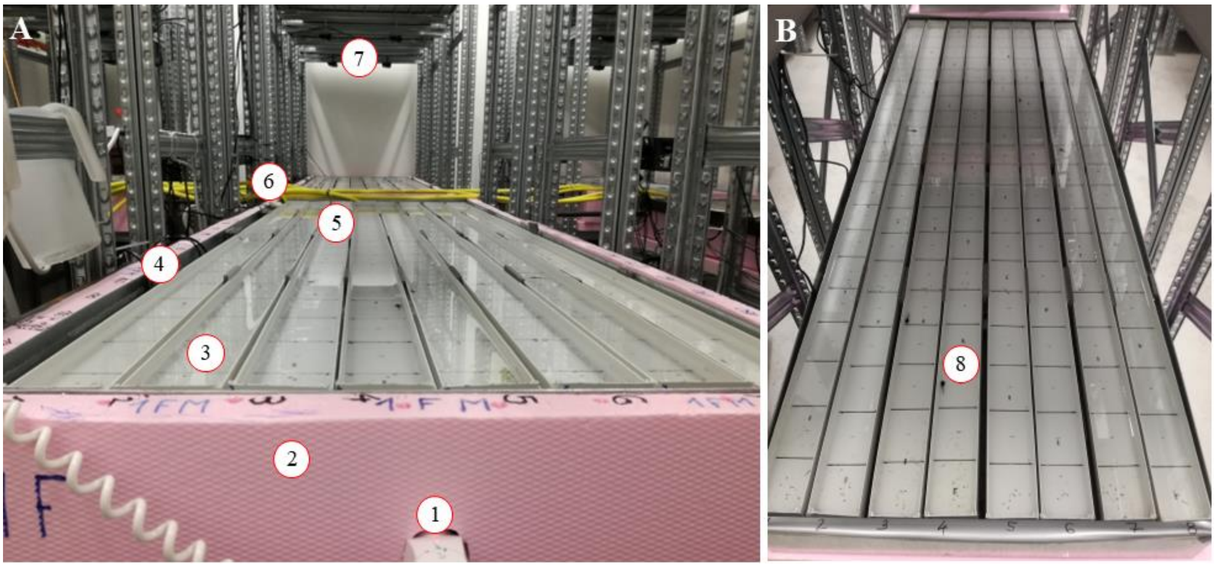

**Supplementary Figure S3**

Temperatures measured along half-pipes in the two thermal treatments when no tadpoles were present. Boxplots show the median (black dot) and interquartile range (box), whiskers extend to the most extreme data points within 1.5 × interquartile range from the box; values more extreme than that are represented by empty circles. For each boxplot, the sample size is 160 measurements (each one of 40 half-pipes measured 4 times).

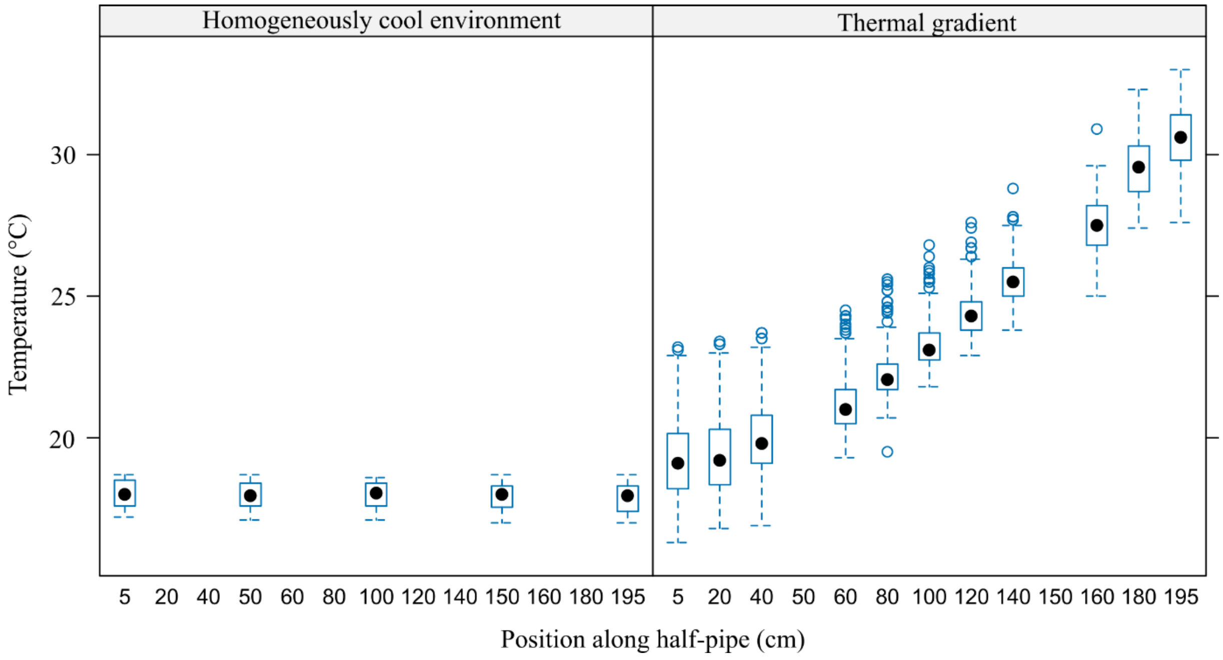
